# Metazoan Diversity in Chilean Hypersaline Lakes Unveiled by Environmental DNA

**DOI:** 10.1101/2024.10.01.616191

**Authors:** Mattia Saccò, Matthew A. Campbell, Pablo Aguilar, Gonzalo Salazar, Tina E. Berry, Matthew J. Heydenrych, Angus Lawrie, Nicole E. White, Chris Harrod, Morten E. Allentoft

**Affiliations:** Subterranean Research and Groundwater Ecology (SuRGE) Group, Trace and Environmental DNA (TrEnD) Lab, School of Molecular and Life Sciences, Curtin University, Perth, Western Australia, Australia; Department of Chemistry, Life Sciences and Environmental Sustainability, University of Parma, Parma, Italy; Trace and Environmental DNA (TrEnD) Lab, School of Molecular and Life Sciences, Curtin University, Perth, Western Australia, Australia; Laboratorio de Complejidad Microbiana, Instituto Antofagasta and Centro de Bioingeniería y Biotecnología (CeBiB), Universidad de Antofagasta, Antofagasta, Chile; Departamento de Biotecnología, Facultad de Ciencias del Mar y Recursos Biológicos, Universidad de Antofagasta, Antofagasta, Chile; Millennium Nucleus of Austral Invasive Salmonids - INVASAL, Concepción, Chile; eDNA Frontiers, Curtin University, Bentley, Western Australia, Australia; School of Biological Sciences, University of Western Australia, Crawley, Western Australia, Australia; Instituto de Ciencias Naturales Alexander von Humboldt, Universidad de Antofagasta, Avenida Angamos 601, Antofagasta, Chile; Lundbeck Foundation GeoGenetics Centre, Globe Institute, University of Copenhagen, Copenhagen, Denmark

**Author notes:** **Correspondence:** Mattia Saccò.

**Keywords:** Environmental DNA, Hypersaline lakes, Metazoan biomonitoring, Biodiversity, Conservation

## Abstract

Saline and hypersaline wetlands are biodiversity hotspots for metazoans such as aquatic invertebrates and wading birds. However, the survival of these habitats and their biota is increasingly threatened by a combination of pressures from climate change and extractive processes, jeopardizing the long-term ecological functioning of these ecosystems. With the goal of improving conservation efforts through state-of-the-art survey techniques in hypersaline ecosystems, this study tests the use of environmental DNA (eDNA) methods for metazoan biomonitoring. We employed a multi-assay approach utilizing three genetic markers—12S rRNA, 18S rRNA, and COI —to analyze biodiversity in two types of environmental substrates, sediment and water. These samples were collected from three hypersaline lakes situated at high altitude in Northern Chile: Salar de Atacama (Laguna Puilar), Salar de Pujsa, and Salar de Tara. In addition, we compared the eDNA outputs with results generated from aquatic macroinvertebrate assessments using kick-nets to evaluate the potential for complementary sampling approaches. Our eDNA analyses revealed a total of 21 and 22 taxa across the three hypersaline lakes in sediment and water, respectively. Within both substrates, the highest diversity was found in Salar de Tara (15 taxa within sediment and 13 taxa from water). Our multi-assay design was able to detect a range of resident hypersaline taxa with different conservation status, spanning from rotifers (*Encentrum*) to endangered snails (*Heleobia atacamensis*), to amphipods (*Hyalella*) and flamingos (*Pheonicopterus*). Macroinvertebrate presence/absence data derived from conventional kick-net surveys further validated Salar de Tara as the most biodiverse system. Compared to net-based assessments, eDNA analysis allowed more refined taxonomic assignments for copepods and ostracods, while certain taxa such as Ephydridae or Hirudinea were not detected through molecular tests. Overall, this study provides evidence that eDNA is an effective tool to elucidate fine scale taxa assemblages and can refine conservation efforts in hypersaline lakes. Given the fast pace of research developments in the field of molecular ecology, eDNA hosts great potential to become a central actor in hypersaline bioassessments in the near future.

## 1 Introduction

Saline and hypersaline lakes worldwide provide immense ecological value (e.g., carbon sinks, microbial diversity hotspots) and host an array of metazoan communities, including charismatic birds such as flamingos and functionally pivotal groups of macroinvertebrate fauna (Balushkina et al., 2009; Hammer, 1986; Saccò et al., 2021b). For instance, the largest saltwater lake (4,400 km^2^) in the western hemisphere – the Great Salt Lake (Utah, USA), supports 338 migratory bird species which depends on the resident benthic invertebrates as an essential primary food source for their migrations along the Pacific Flyway (Baxter and Butler, 2020). In the southern hemisphere, the Salar de Atacama (3,000 km^2^), the third-largest saline pan in the world after Salar de Uyuni in Bolivia (10,582 km^2^) and Salinas Grandes in Argentina (6,000 km^2^), hosts one vulnerable (*Phoenicoparrus andinus*) and two near threatened (*Phoenicopterus chilensis* and *Phoenicoparrus jamesi*) species of flamingos (Gajardo and Redón, 2019). Located in the pre-Andean depression of the Atacama Desert together with more than fifty other hypersaline lakes, Salar de Atacama has been the focus of extensive geohydrological studies (e.g., (Finstad et al., 2016; Lowenstein et al., 2003; Marazuela et al., 2019), although with far less work being generated on ecological patterns.

In the Atacama region and worldwide, molecular-based studies on saline and hypersaline lakes have predominantly focused on microbes rather than metazoans. Microbial processes in these systems are essential to biogeochemical flows and ecosystem functioning, including nutrient cycling, carbon fixation, and the degradation of organic matter (Oren, 2015; Oren et al., 2009). This microbial focus has attracted significant industrial interest in regions with strong extractive drives due to the mineral resources found in these lakes (Garcés and Álvarez, 2020; Kidder et al., 2020). Consequently, there has been a diminished interest in studying macro-organismal groups. This trend is likely exacerbated by the lack of methodological and technological advancements, despite the important role metazoans play in energy transfer and habitat modification. As a result, our understanding of the functional dynamics regulating these ecosystems and the services they provide is limited. The generally low diversity and visibility of metazoans, combined with the significant logistical and sampling challenges of studying larger organisms, further diverts research attention away from metazoan taxa such as zooplankton, invertebrates, and birds.

Investigating metazoans in saline ecosystems is critical due to their role as bioindicators of environmental health and their unique biodiversity, which can reveal novel adaptations and evolutionary processes (Zadereev et al., 2020). Recent advancements in DNA sequencing technologies, bioinformatics, and sampling methodologies are facilitating a more comprehensive exploration of biodiversity, particularly within unique environments such as hypersaline lakes or saltpans (Saccò et al., 2021b). One non-invasive technique gaining considerable momentum in aquatic biomonitoring studies is environmental DNA (eDNA) analysis – the detection of the total pool of DNA isolated from environmental samples ((Takahashi et al., 2023) and references therein). eDNA analysis offers several logistical benefits over traditional methods (Blackman et al., 2024). Firstly, it enhances efficiency as eDNA sampling requires less time and effort compared to traditional biodiversity surveys, which often involve extensive fieldwork and specimen collection (Effendi et al., 2023). Secondly, eDNA analysis is cost-effective because its non-invasive nature reduces the need for specialized equipment and personnel, lowering the overall cost of biodiversity monitoring (Jo et al., 2022). Lastly, eDNA analysis provides a broader scope by capturing a wide range of species, including those that are elusive or rare, thereby offering a more comprehensive understanding of the ecosystem’s biodiversity (Hansen et al., 2018). These three aspects have the potential to be highly beneficial for hypersaline lake assessments, understudied ecosystems often located in remote poly-extreme environments with potential risks for the health of researchers (due to high altitude and extreme temperature ranges, strong UV radiation, etc.) when conventional survey methods (e.g., trapping, counts) are utilised. However, eDNA applications in inland saline ecology to date are scarce, mainly due to the technical challenges with high salt concentration and elevated UV radiation affecting the DNA extraction processes and eDNA persistence/quality in the environment, respectively (e.g., (Saccò et al., 2021b; Tazi et al., 2014).

As demonstrated in other remote and extreme environments (such as deep oceans or groundwaters), eDNA has the potential to become a routinary tool for assessing biodiversity patterns due to recent analytical and technological progress (Hinz et al., 2022; Saccò et al., 2022; Takahashi et al., 2023). Building upon successful eDNA metabarcoding assessments of invertebrate community patterns in Australian hypersaline lakes (Campbell et al., 2023), this study aims to evaluate the applicability of eDNA metabarcoding techniques in delineating broader metazoan communities present in both water and sediment samples from hypersaline lakes in Chile. Specifically, we aim to (i) use an eDNA approach to describe the avian and invertebrate biodiversity present in three hypersaline lakes located at different altitudes in Northern Chile and to (ii) compare the macroinvertebrate biodiversity detected by eDNA against conventional net sampling and provide conservational guidelines. The results of this study bring new light on hypersaline diversity patterns and provide preliminary evidence on the validity and applicability of the eDNA technique in inland high salt conditions.

## 2 Materials and methods

### 2.1 Study area

Field work was carried out at three hypersaline lakes located in the Los Flamencos National Reserve, which comprises a total of seven sectors from the Salar de Atacama to the salares and lagoons of the altiplano in the Antofagasta region, covering an area of 73,986.5 ha (CONAF, 2008). Water, sediment and macroinvertebrate samples were collected in October 2021 from three sites: Salar de Tara (4322 masl), Salar de Pujsa (4514 masl) and Laguna Puilar (2306 masl). Samples were collected from three sampling points per each salar to encompass for the environmental heterogeneity of the sites (Figure 1).

**Figure 1.**
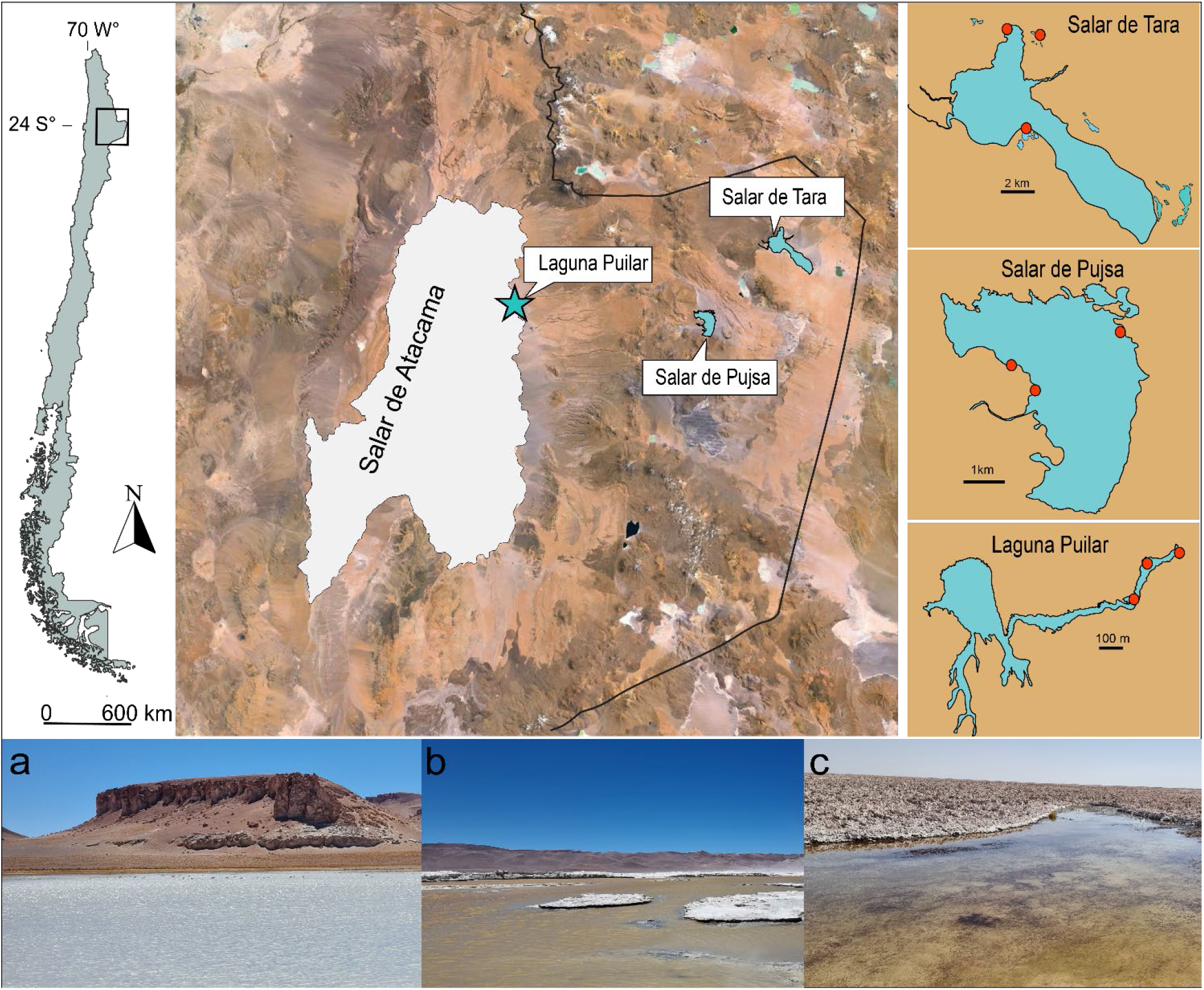
Location of sampling points at laguna Puilar (Salar de Atacama), Salar de Pujsa and Salar de Tara within the Los Flamencos National Reserve. The map depicts the difference in size of each salar and the three sampling points (in red) where samples were collected (Laguna Puilar displayed not in scale within the Salar the Atacama for representative reasons). At the bottom of the figure, some photos of the landscapes are shown: a) Salar de Tara, b) Salar de Pujsa and c) Salar de Atacama (Laguna Puilar).

### 2.2 Kick-net sampling for macroinvertebrate diversity

Benthic invertebrate diversity was estimated using kick net sampling (mesh = 500 μm), a methodology that is frequently used for qualitative and semi-quantitative assessments in rivers and streams (Everall et al., 2017), but also in hypersaline lakes (Herbst, 1988). All macroinvertebrates found in the kick net were passed through a sieving system, consisting of four sieves of decreasing mesh size (500, 200, 100 and 44 μm) to remove excess sediment and eventually trap meiofauna (invertebrates smaller than 500 μm) from the sediments themselves. Samples were stored in plastic bags and then analysed in the laboratory with a binocular stereo microscope. Taxa were identified to the lowest level possible by using taxonomic keys (e.g., (Domínguez and Fernández, 2009; González, 2003; Roig-Juñent et al., 2014) and a matrix with the diversity data was generated.

### 2.3 eDNA sample collection

Water samples were collected using a 3 L sterilized bottle, and filtration was conducted *in situ* using a vacuum pump equipped with 0.4 and 0.22 µm mesh size filters. These filters were preserved in RNAlater within 2 mL tubes and stored at -20 °C in darkness to maintain integrity. Although the initial goal was to obtain one 0.44 µm and one 0.22 µm filter for each sample, clogging issues due to the high salt concentrations in water occasionally necessitated the use of additional filters. For representative purposes, filters were combined to homogenise the volume analysed per sampling point (for both the 0.44 and the 0.22 µm mesh size filters) which finally ranged between 200 to 237 ml (Supplementary Table 1). Additionally, one negative control for each sampling point, consisting of filtering 1 L of distilled water in the field, was included in the analysis. Sediment samples from the top 3 cm were collected in triplicate at each sampling point. These were stored in sterile 50 ml tubes containing analytical grade ethanol (99.9 %) and kept frozen at –20 ºC until further analysis.

### 2.4 eDNA extraction and purification

All laboratory processes were conducted in dedicated laboratories within the Trace and Environmental DNA (TrEnD) Laboratory, Curtin University, Perth, Western Australia, and all controls were processed alongside samples through to sequencing. In total, 31 filters were used for the samples and 18 for the negative controls. DNA from the water was extracted using a modified Qiagen DNeasy blood and tissue kit protocol (Qiagen, Germany). The proportion of the filter papers from each sample replicate that provided the equivalent of approximately 225 mL of filtered water was digested (see Supplementary Table 1 for actual amounts). This quantity was chosen because it represented the minimum amount of water filtered at any one site. As all or a portion of some filters were digested for extraction, all of each of the sample control filters were digested alongside the sample filters. Two extraction controls were also included. Samples were digested using 810 µL Buffer ATL and 120 µL Proteinase K and were incubated overnight at 56 °C with rotation. The following day 400 µL of the digest was combined with 400 µL Buffer AL and 400 µL 100 % Ethanol. The remainder of the protocol was as prescribed by Qiagen with the addition of a second wash with Buffer AW2 to remove any extra salts introduced by the increased ratio of Proteinase K. The final modification was elution with a double elution (2 × 50 µL) to maximise DNA recovery for a total of 100 µL in Buffer AE.

Sediment samples were extracted using a modified Power Soil Pro Kit protocol (Qiagen, Germany). Samples were defrosted and homogenised with two to four 10 mm silver balls were added to each falcon tubes (depending on the volume) and placed in the TissueLyser (Qiagen, Germany) for 30 s at 30 Hz. Approximately 250 mg of the slurry was subsampled into Powerbead Tubes. Four small, sterile glass beads were also added to all sample tubes and two extraction controls. After the addition of 800 µL of Buffer CD1 the samples were placed in the TissueLyser twice at 25 Hz for 5 min. Extractions were continued using a (QIAcube Qiagen, Germany) and following the Power Soil Pro IRT method. Extractions were eluted in a final volume of 100µL AE buffer.

### 2.5 eDNA metabarcoding

The eDNA in the samples was analysed using three metabarcoding assays: 12S rRNA (L1091/H1985 - Cooper et al., 1994), 18S rRNA (18S_1F/18S_400R - Pochon et al., 2013), and COI (fwhF2/fwhR2n - Vamos et al., 2017) (Supplementary Table 2). To evaluate DNA extraction inhibition and to determine the required dilution for optimal amplification, PCR reactions were performed on each DNA extract by adding DNA template directly to the PCR master mix (neat), then performing serial dilutions (Water to 1 in 10, and Sediment to 1 in 10 and to 1 in 100). The PCRs were performed at a final volume of 25 µL where each reaction comprised of: 1× PCR Gold Buffer (Applied Biosystems), 0.25 mM dNTP mix (Astral Scientific, Australia), 2 mM MgCl_2_ (Applied Biosystems), 1 U AmpliTaq Gold DNA polymerase (Applied Biosystems), 0.4 mg/mL bovine serum albumin (Fisher Biotec), 0.4 µM forward and reverse primers, 0.6 μl of a 1:10,000 solution of SYBR Green dye (Life Technologies), and 2 µL template DNA or dilution. PCRs were performed on StepOne Plus instruments (Applied Biosystems) with the following cycling conditions: 95 °C for 5min, followed by 50 cycles of: 95 °C for 30 s, Assay Tm °C (Supplementary Table 2) for 30 s, 72 °C for 45 s, then a melt-curve analysis of: 95 °C for 15 s, 60 °C for 1 min, 95 °C for 15 s, finishing with a final extension stage at 72 °C for 10 min.

After selection and creation of the optimal dilution (neat, 1 in 10, or 1 in 100), PCRs were repeated in duplicate as described above and instead using unique, single use combinations of 8 bp multiplex identifier-tagged (MID-tag) primers as described in Koziol et al. (2019) and van der Heyde et al. (2020). Master mixes were prepared using a QIAgility instrument (Qiagen) in an ultra-clean lab facility, with negative and positive PCR controls included on every plate. A sequencing library was created by combining samples into equimolar mini-pools based on the PCR amplification results from each sample. The mini-pools were then analysed using a QIAxcel fragment analyser (Qiagen) to combine them into equimolar concentrations and form the final sequencing libraries. Each library was then size selected (Pippin Prep: Sage Sciences) with 2 % dye-free cassettes, cleaned using a QIAquick PCR purification kit, quantified by Qubit (Thermo Fisher), and diluted to 2 nM. All libraries were sequenced on an Illumina MiSeq instrument using MiSeq V2 sequencing kits with custom sequencing primers.

### 2.6 Bioinformatics and data analysis

Raw metabarcoding sequence data was filtered and analysed using the eDNAFlow pipeline (Mousavi-Derazmahalleh et al., 2021) with custom filtering parameters applied (--minAlignLeng ‘12’, --minLen ‘(see Supplementary table 2)’, --minsize ‘2’, perc_identity ‘95’ (‘90’ for 18S), --qcov ‘100’, -- minMatch_lulu ‘97’). Generated ZOTUs were queried against the nucleotide database NCBI (GenBank) and assigned to the species level where possible or dropped back to the lowest common ancestor if multiple possible taxonomic assignments were given. Low abundant ZOTUs without at least one sample above 0.003% sequences assigned, were considered unreliable and excluded from the dataset (Vamos et al., 2017). Any ZOTUs from field, extraction and negative controls (a total of 1 field control per water sampling site, 4 extraction controls and 1 negative controls per assay) were removed from the dataset, and 3 different positive controls (one per assay) were also sequenced for testing purposes. To depict the diversity of taxa identified, a phylogenetic tree was constructed utilizing phyloT (Phylogenetic Tree Generator) relying on the taxonomy from the National Center for Biotechnology Information (NCBI), and subsequently rendered using the Interactive Tree of Life (iTOL) (Letunic and Bork, 2021).

## 3 Results

### Biodiversity of the Chilean precordillera

Following read filtering, an average of 19,411 reads for 12S rRNA, 75,662 reads for 18S rRNA, and 4,858 reads for COI were achieved per sample. Across the three wetlands and two substrates sampled, 31 taxa were found via eDNA approaches. Salar de Tara exhibited the highest diversity in both sediment (15 taxa) and water (13 taxa), followed by Laguna Puilar (6 taxa in sediment and 11 taxa in water) and Salar de Pujsa (1 taxon in sediment and 4 taxa in water). The richness of taxa was comparable between water (n = 21) and sediment (n = 22) samples. For visual representation, see Figure 2 displaying the taxa distribution. Several groups were only recorded from sediments (*Epirodrilus*, Branchinectidae, *Cyprideis, Limnocythere*, Orthocladiinae, *Trhypochthonius*, Cyatholaimidae, *Gieysztoria* and *Encentrum*) or in water (*Nais, Hyalella, Liposcelis, Smittia, Chroicocephalus*, Scolopacidae, *Spatula, Heleobia atacamensis, Macrostomum* and *Jakoba libera*). The only taxon that was found in all the hypersaline lakes was the Greater Flamingo (*Phoenicopterus*). Arthropoda was the most diverse Phylum with 12 taxa within Hexanauplia (5 taxa), Ostracoda (4 taxa) and Insecta (3 taxa) followed by Chordata with Aves (6 taxa), while several other Phyla yielded one taxon (e.g., Tardigrada and Malacostraca).

**Figure 2.**
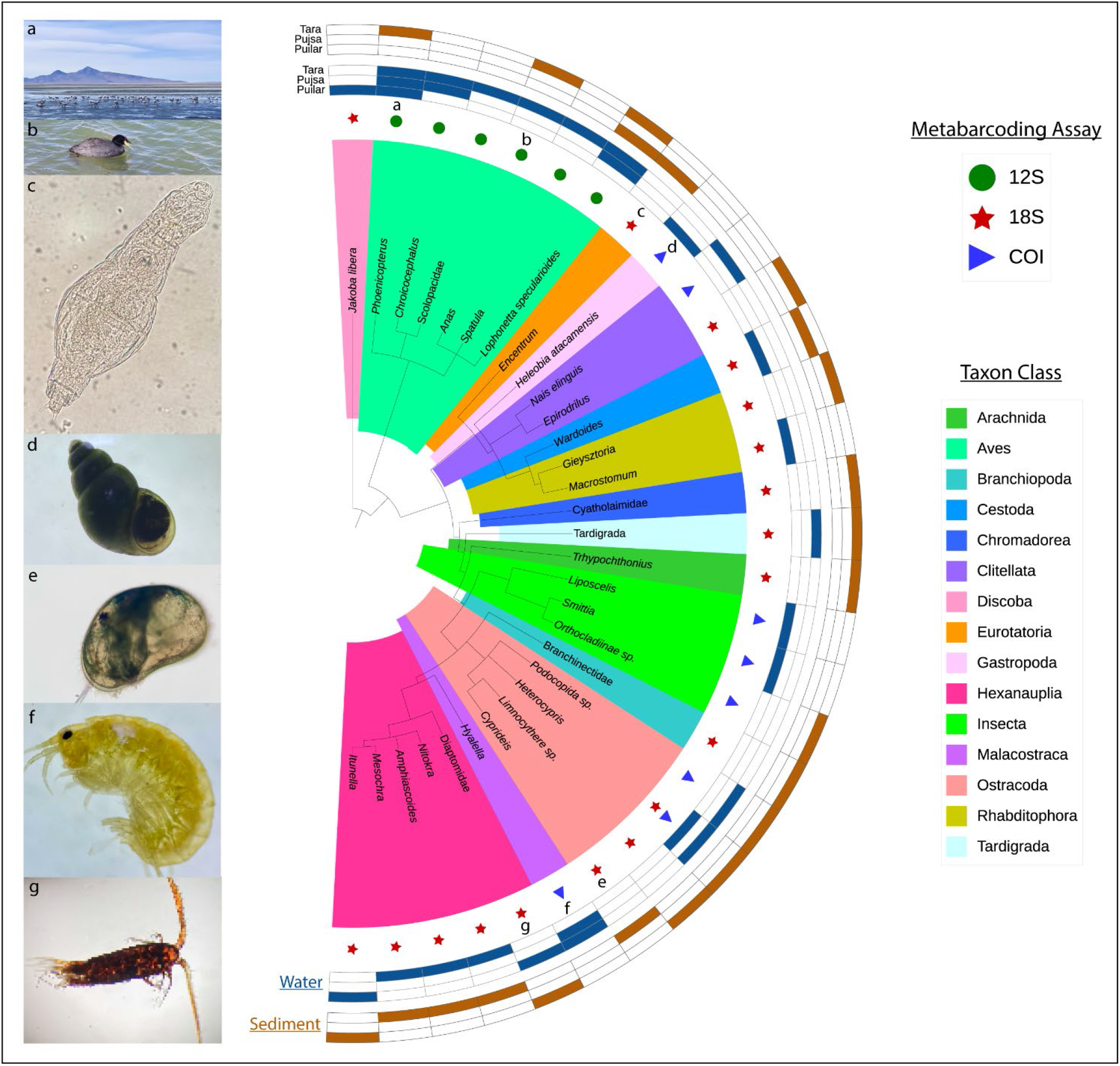
iTOL graph showing the taxa found at the three hypersaline lakes (salar de Tara, laguna Puilar (salar de Atacama) and salar de Pujsa) in water and sediment through the multi-assay (12S, 18S and COI) approach. On the left photos of some specimens from the field and sorting in the laboratory are shown.

Metabarcoding analysis using the bird universal primer 12S yielded 6 bird species. The 18S dataset detected 18 taxa across a broader taxonomic spectrum and notably captured copepod diversity consistently across both water and sediment samples from each salar surveyed. Additionally, the 18S assay proved effective in detecting other microscopic taxa such as cestodes (Wardoides) and rotifers (Encentrum). Conversely, COI performed better than the other two assays at detecting gastropods (*Heleobia atacamensis*), dipterans (e.g., *Smittia*) and amphipods (*Hyalella*).

### eDNA vs net sampling

Taxonomic assessment with eDNA and net sampling techniques revealed distinct patterns in the distribution and detection of macroinvertebrate taxa across the hypersaline lakes (Table 1). Within Clitellata, Hirudinea was exclusively identified in Salar de Tara through net sampling. Oligochaeta, specifically within the family Naididae, was detected in Laguna Puilar using net sampling methods. At a finer taxonomic resolution, the species *Nais elinguis* was discerned in Salar de Tara through metabarcoding techniques (COI). Interestingly, Naididae was exclusively detected via net sampling in Laguna Puilar, contrasting with the specific identification of *Nais elinguis* through eDNA metabarcoding in Salar de Tara. Among Gastropoda, *Heleobia* was identified in Laguna Puilar using both methodologies, with species-level identification achieved for *Heleobia atacamensis* through metabarcoding (COI). Crustaceans, notably Amphipoda such as *Hyalella sp*., were consistently found across all three ecosystems using both net sampling and metabarcoding approaches, although *Hyalella sp*. eluded detection via eDNA analysis at Laguna Puilar. The Copepoda community included Calanoida (*Boeckella sp*.) observed in Salar de Tara via net sampling, Diaptomidae in Pujsa through metabarcoding, and Cyclopoida in Tara via metabarcoding. Harpacticoida species were observed in Tara and Puilar through net sampling, and in Salar de Pujsa through metabarcoding, with various copepod taxa (*Nitokra* sp., *Amphiascoides* sp., *Mesochra* sp., *Itunella* sp.) identified across multiple ecosystems using metabarcoding techniques. Ostracoda species such as *Podocopida* and *Heterocypris* sp. were detected in Salar de Pujsa and Salar de Atacama (Puilar) through metabarcoding, respectively, while Branchiopoda (Chydoridae and Ilyocryptidae) were detected in Salar de Tara through net sampling. Insecta included Hemiptera (Corixidae) found in Salar de Tara via net sampling, and Diptera families Ephydridae, Dolichopodidae, and Chironomidae identified using both sampling methods. *Smittia* sp. was detected in Laguna Puilar through metabarcoding.

**Table 1.**
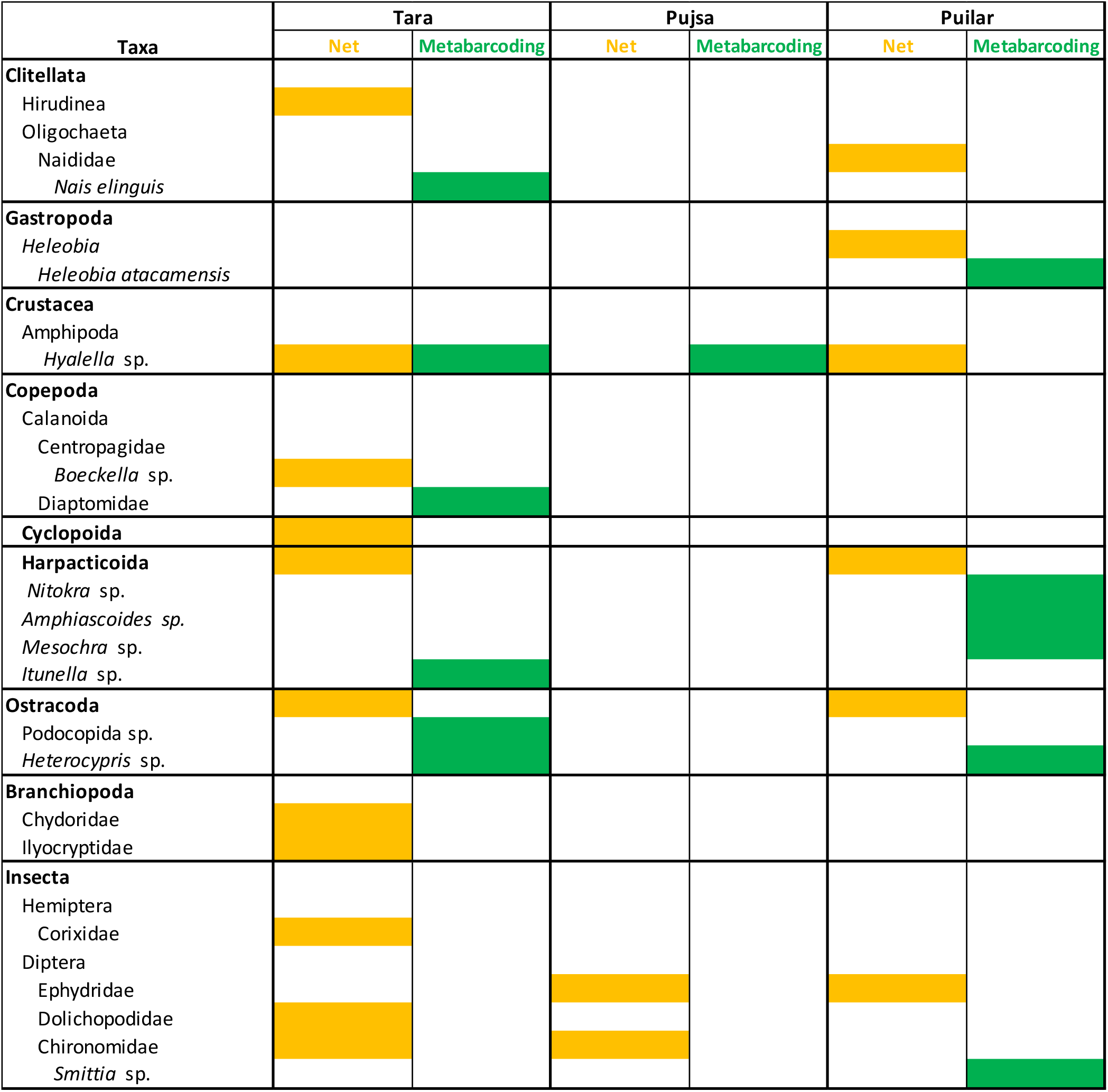
Comparison of morphology-based taxonomic data obtained via kick-net surveying versus results through eDNA metabarcoding analysis for aquatic macroinvertebrates within the three hypersaline lakes targeted (Salar de Tara, Salar de Pujsa and Laguna Puilar (Salar de Atacama)). Positive detected indicated with orange (net) and green (metabarcoding).

## 4 Discussion

This study provides the first multi-assay eDNA analysis in water and sediment samples from Chilean high altitude hypersaline lakes. We successfully generated eDNA metabarcoding sequence data from both water and sediment samples, overcoming the inherent challenges associated with DNA preservation in these environments, which include high salinity, UV exposure, and difficulties in filtering saline water to collect eDNA. The findings from the eDNA assays revealed several endemic taxa (i.e., *Heleobia atacamensis, Pheonicopterus*) frequently reported in the hypersaline lakes of the Chilean Precordillera and Altiplano. These results extend recent studies indicating that eDNA approaches represent a promising and effective method for assessing the biodiversity of salt lakes (Campbell et al., 2023; Saccò et al., 2021a). All the taxa, apart from the ostracod *Heterocypris*, were only identified by a single assay, confirming the need for the implementation of multiple universal assays in eDNA studies attempting to examine metazoan community composition (Takahashi et al., 2023). At ecosystem level, our integrated eDNA-based findings demonstrate that the Salar de Tara hosts the highest metazoan diversity. These results underscore the necessity for enhanced biomonitoring activities (involving ideally extensive sampling campaigns over multiple seasons) to fully disclose the biodiversity patterns on site and maintain the ecohydrological integrity of the wetland (García-Sanz et al., 2021).

### Avifaunal diversity

Six different waterbird taxa were identified in the eDNA data across three sites with Salar de Tara supporting all of them. Out of the three endemic species breeding in the Atacama Desert and Chilean Altiplano (*Phoenicopterus chilensis, Phoenicoparrus jamesi* and *Phoenicoparrus andinus*), only Chilean Flamingos (*Phoenicopterus* sp.) - the most widespread and abundant overall (Gutiérrez et al., 2022 and references therein) - were identified in waters from all three sites. The reliable detection of Chilean Flamingos using eDNA from water samples is important from a conservation perspective because flamingos are useful “flagship species” to promote the conservation of hypersaline lagoons which are under threat (Gajardo and Redón, 2019; Williams, 1998). In addition, Chilean Flamingos are known for carrying long migratory routes and for making use of several hypersaline habitats for feeding and breeding (Polla et al., 2018), and eDNA can help detect recent presence in the environment.

On the other side, the smaller *Phoenicoparrus* specimens were abundant in all the three hypersaline lakes (personal observation Gonzalo Salazar, Figure 3), but the two species (*Phoenicoparrus jamesi* and *Phoenicoparrus andinus*) belonging to this genus were not detected *via* eDNA approaches. Stochastic sampling factors and reference sequence availability for the genus *Phoenicoparrus* (e.g., no 12S sequences for this genus are currently openly available on GenBank) may have influenced the non-detection, stressing the need for development of species-specific assays that could even dig into gender identification of specimens as indicated by (Chapman, 2012).

**Figure 3.**
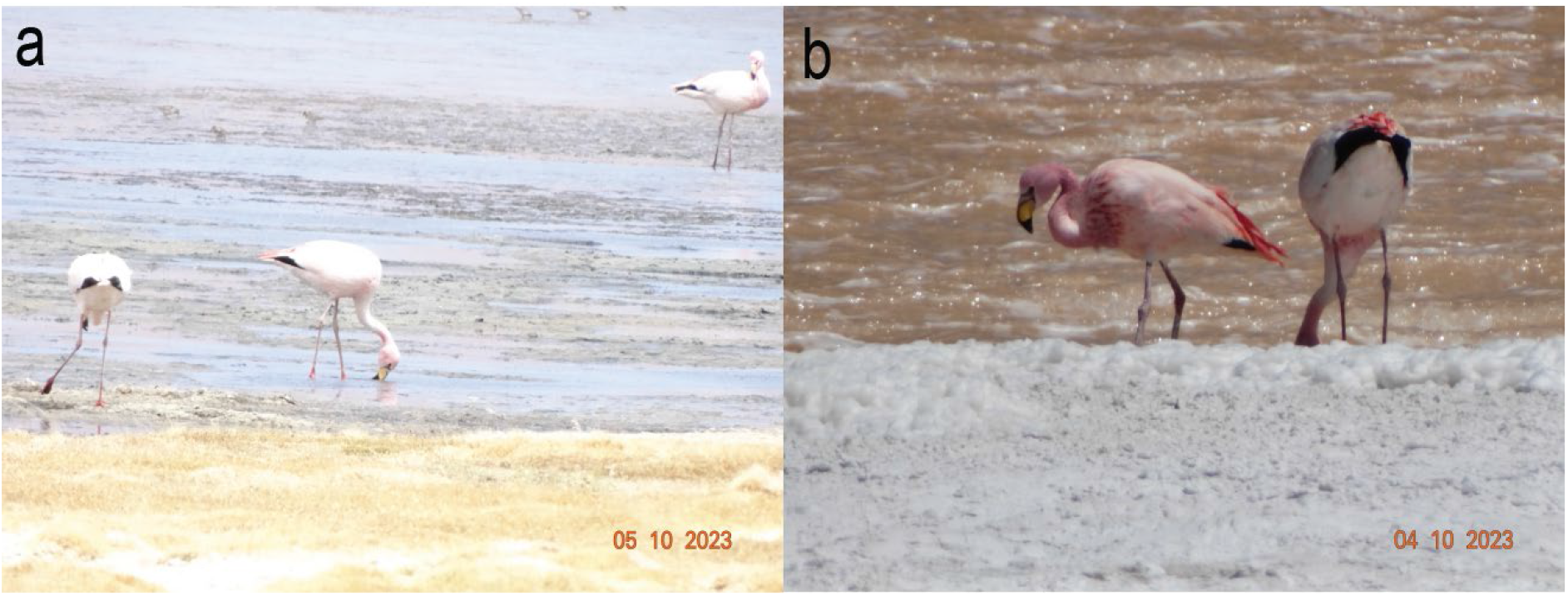
Photos of *Phoenicoparrus Jamesi* individuals collected during fieldwork at Salar de Tara (a) and Salar de Pujsa (b). Credit: Gonzalo Salazar

Another taxon of wading birds identified through aquatic eDNA is the South American crested duck (*Lophonetta specularioides*), whose distribution ranges in Chile include the Altiplano, a high-altitude plateau area situated among the high peaks of the Andes at over 4000 m a.s.l. (Muñoz-Pedreros et al., 2015). Our molecular analyses revealed the presence of this species of native South American duck in waters of the Salar de Pujsa and Salar de Tara, the two wetlands located within the puna. Other avifaunal groups identified through aquatic eDNA and frequently reported in the region were gulls (*Chroicocephalus*) and sandpipers (Scalapacidae). Only two taxa were found in sediment (*Phoenicopterus* and *Anas* and both at Salar de Tara), suggesting that this substrate is not ideal for investigating the use of habitat of avifaunal species. Overall, these findings indicate that eDNA research has potential to build on classic visual avifaunal surveys in hypersaline lakes, as already proved in waterholes wetlands (e.g., (Li et al., 2023) and for rare and secretive mash bird species (e.g., (Neice and Mcrae, 2021).

### Invertebrate diversity

The majority of taxa identified in the waters and sediments of the three hypersaline lakes *via* eDNA approaches (n = 25) belonged to invertebrate groups. In sediment samples, most taxa were benthic, including nematodes (Cyatholaimidae), rotifers (*Encentrum*), oligochaetes (*Epirodrilus*), chironomids (Orthocladiinae), platyhelminthes (*Gieysztoria*) or potentially derived from dead tissue of nektonic species (ostracods *Cyprodeis* and *Limnocythere*). In water samples, our metabarcoding analyses identified the critically endangered gastropod species *Heleobia atacamensis* that is endemic to the Salar de Atacama. This snail was originally described in Tilopozo, South of the Salar de Atacama in 1860 (Collado et al., 2013). However, recent phylogenetic analyses and multi-locus research confirm that a much greater population exists (Collado et al., 2023) and reference therein), with specimens found at Peine, Tilomonte and Puilar wetlands, amongst others. Our results confirm the presence of the species at Laguna Puilar, providing evidence of the validity of eDNA approaches for hypersaline biomonitoring when reference sequence data is available. Our molecularly based design was also successful in bringing new light to the diversity of aquatic micro-animals (e.g., rotifers, tardigrades), amphipods (*Hyalella*), and certain meiofaunal groups such as copepods (harpacticoids at Laguna Puilar) and ostracods (*Heterocypris* at Tara and Puilar). Further, the eDNA data identified the cestode *Wardoides* that has not been reported in hypersaline lakes previously highlighting the potential capacity for eDNA methods to reveal otherwise overlooked biodiversity in these environments (Beng and Corlett, 2020). The only calanoid detected through molecular approaches was in Salar de Tara and was only able to be identified to the family Diaptomidae, a widespread and specious family comprising species that have adapted to a range of different aquatic environments including inland saline lakes (Bayly, 1993), but not reported in Northern Chile to date. Net surveys from the same salar identified the presence of the calanoid copepod genus *Boeckella* sp., a taxon that is known to occur in salt lakes form the Chilean Altiplano (Williams, 1998).

Overall, the eDNA approaches in our study generated greater taxonomic resolution compared to morphological identifications from net sampling in most faunal groups, particularly aquatic micro-animals (e.g., rotifers, tardigrades, copepods and ostracods) (Table 1). Nonetheless, the designed eDNA analysis was not able to detect some macroinvertebrate groups found with planktonic nets such as Ephydridae, Cladocera or Hirudinea, also with other groups being not recorded at sites when present (*Hyalella* at laguna Puilar, Salazar’s personal communication). This aspect stresses the need for eDNA-net complementary approaches, and the development of reference DNA databases for hypersaline lakes to avoid false-negative results due to a lack of reference sequences or primer biases (Takahashi et al., 2023).

Indeed, the absence of the brine shrimps *Artemia* (Artemiidae) in this study is also noteworthy, as this taxon is considered a conspicuous component of southern South American salt lakes (Williams, 1998). Neither eDNA nor kick-net samples nor samples suggested the presence of *Artemia* in the three lakes studied, although an Anostracan of the family Branchinectidae was detected in sediment samples at Salar de Tara. These results could indicate that *Artemia* does not provide a major food source for flamingos in the three lakes targeted in this study, in line with Polla et al., (2018). Previous research also indicate that famingos’ diet preferences are diverse and vary depending on seasonal food availability and species-specific trophic behaviours (Dorador et al., 2018; Gajardo and Redón, 2019), an eco-evolutionary pattern eventually contributing to the coexistence of sympatric species in similar waters (Polla et al., 2018). Nonetheless, given the relatively limited scope of this study, other stochastic factors such as sampling random effects or PCR primer binding issues might have played a role in shaping these findings presented, and further research will be necessary to confirm these trends.

### Conservational considerations

Our findings show that species of conservation concern (e.g., Chilean Flamingos and *H. atacamensis*) are utilising salt lakes in the Chilean precordillera and Altiplano. This is crucial to recognise as saline lakes are increasingly being exposed to mining exploration for minerals such as lithium (Gajardo and Redón, 2019). A recent study targeting the Chilean portion of the so called “Lithium Triangle” (Salar de Atacama, Salar de Tara and Salar de Pujsa) by (Gutiérrez et al., 2022) provided evidence of decreased flamingo diversity in correspondence of lakes subjected to increased lithium extractions over the last 30 years. This trend is likely to be indicative of overall ecological impoverishment of Salares, an alarming dynamic that requires urgent targeted conservational management strategies to avoid environmental collapse (García-Sanz et al., 2021). Worldwide, the combination of uncontrolled resource extraction and climate change effects is threatening the long-term ecological integrity of hypersaline lakes. Inter-disciplinary approaches are the gateway to investigate such complex dynamics, an essential task in a world that is unequivocally going to face increased freshwater salinization due to anthropogenic impacts (Cunillera-Montcusí et al., 2022). Indeed, eDNA provides a powerful complementary platform for unveiling otherwise overlooked biodiversity patterns under high-salt conditions.

## 5 Conflict of Interest

The authors declare that the research was conducted in the absence of any commercial or financial relationships that could be construed as a potential conflict of interest.

## 6 Author Contributions

M.S.: conceptualization, methodology, validation, formal analysis, investigation, writing – original draft, project administration; M.C.: investigation, curation, formal analysis, writing – review & editing; P.A.: sampling, investigation, writing – review & editing; T.B.: investigation, writing – review & editing, M.J.H.: investigation, writing – review & editing; A.L.: writing – review & editing; N.E.W.: writing – review & editing; G.S: sampling, investigation; writing – original draft; writing – review & editing; C.H: sampling, writing – review & editing; M.E.A: writing – review & editing, project administration.

## 7 Funding

This research forms part of the eDNA for Global Biodiversity (eDGES) program. P.A, G.S., and C.H. received funding from ANID–Millennium Science Initiative Program–NCN2021-056.

## 8 Acknowledgments

This work was supported by resources provided by the Pawsey Supercomputing Centre with funding from the Australian Government and the Government of Western Australia. M. S., G. S. and N. E. W acknowledge support from the BHP-Curtin alliance within the framework of the “eDNA for Global Environment Studies (eDGES)” program.

## 9 Data Availability Statement

All additional data are available in the Supplementary material and the DNA-based data are archived in the Dryad repository (https://doi.org/10.5061/dryad.80gb5mm08).

## Notes

### Competing Interest Statement

The authors have declared no competing interest.

